# Developing T cells form an immunological synapse for passage through the β−selection checkpoint

**DOI:** 10.1101/732511

**Authors:** Amr H. Allam, Mirren Charnley, Kim Pham, Sarah M. Russell

## Abstract

The β-selection checkpoint of T cell development tests whether the cell has recombined its genomic DNA to produce a functional T Cell Receptor β (TCRβ) receptor. Passage through the β-selection checkpoint requires the nascent TCRβ protein to mediate signaling through a pre-TCR complex. In this study, we show that developing T cells at the β-selection checkpoint establish an immunological synapse in *in vitro* & *in situ,* resembling that of the mature T cell. The immunological synapse is dependent on two key signaling pathways known to be critical for the transition beyond the β-selection checkpoint, Notch and CXCR4 signaling. *In vitro* and *in situ* analyses indicate that the immunological synapse promotes passage through the β-selection checkpoint. Collectively, these data indicate that developing T cells regulate pre-TCR signaling through the formation of an immunological synapse. This signaling platform integrates cues from Notch, CXCR4, and MHC on the thymic stromal cell, to allow transition beyond the β-selection checkpoint.

**Summary:** T cell development requires testing whether genomic rearrangement has produced a T cell receptor capable of transmitting signals. Most T cells fail this test. Here, we show that passage through the β-selection checkpoint requires assembly of a platform to support TCR signaling.

## Introduction

Mature T cells each express a unique T Cell Receptor (TCR) to enable binding and response to specific antigens (Smith-Garvin et al., 2009; Turner et al., 2006). The TCR is created by genomic recombination during T cell development in the thymus (Mallick et al., 1993; Pardoll et al., 1987). To ensure that its TCR is fit for purpose, the developing T cell must survive a series of tests. The first of these tests, termed the β-selection checkpoint, assesses whether the developing T cell has recombined its TCRβ gene appropriately (Carpenter and Bosselut, 2010). Much has been learnt about the signaling and cell fate decisions that depend upon correct TCRβ recombination at this stage (Chann and Russell, 2019). At the β-selection checkpoint, the T cell still lacks a recombined TCRα, but pairs with pTα chain (Groettrup et al., 1993; Raulet et al., 1985). The nascent TCRβ paired with pTα is termed a pre-TCR receptor, and shares many characteristics with the TCR of mature T cells, including association with CD3 and associated cell surface receptors, and triggering of similar signaling cascades (Saint-Ruf et al., 2000; von Boehmer, 2005).

The vast literature on mechanisms of action of the TCR in mature cells has provided a platform for understanding the mechanisms of pre-TCR signaling, but not all characteristics are shared between the two TCR types (Gascoigne et al., 2016). The most profound differences relate to the impact of ligand-binding on higher order organization of signaling. In a mature T cell, interaction of the TCR with antigen presented via Major Histocompatibility Complex (MHC) triggers a highly orchestrated spatial reorganization (Alcover et al., 2016a; Martin-Cofreces et al., 2011). This reorganization involves assembly of a dynamic structure termed the immunological synapse, which enables control over the intensity and duration of the antigen presentation signal during T cell activation and T cell mediated killing of target cells (Dustin et al., 2010; Lee et al., 2003; Lee et al., 2002). A similar structure has not been considered in developing T cells, in part because the lack of a genomically rearranged TCRα chain at the β-selection checkpoint means that the cell cannot undergo canonical binding to peptide-MHC complexes (Mallick et al., 1993). In place of ligand binding and formation of an immunological synapse, models for activation of the pre-TCR initially relied upon the notion that dimerization of TCRβ with pTα stabilizes the protein, paving the way for lipid-raft-based clustering to initiate signaling (Saint-Ruf et al., 2000). However, a number of studies showed that the cysteine residue in the pTα cytoplasmic domain, which is involved in the palmitoylation site, is not required for the DN to DP transition (Aifantis et al., 2002). Moreover, CD3ε dimerization was sufficient for progression beyond β-selection even when there was no raft localization (Levelt et al., 1995; Shinkai et al., 1995). Later studies showed that the pTα extracellular domain undergoes self-oligomerization via four charged amino acids that are required for initiation of the pre-TCR signaling and progression beyond the β-selection (Pang et al., 2010; Yamasaki et al., 2006). On the other hand, signal initiation of the pre-TCR does not require oligomerization, but rather depends on regulating the surface expression of the pre-TCR complexes and their abundance (Mahtani-Patching et al., 2011). Thus, there is not yet a clear consensus as to what triggers productive signaling through the pre-TCR.

It is becoming apparent that signaling through the pre-TCR might be more akin to signaling through the TCR of mature T cells than had previously been considered. Recent findings indicate that the pre-TCR can bind peptide-MHC complexes, and that signaling from the peptide-MHC complex can influence T cell development (Das et al., 2016; Mallis et al., 2015). The affinity of the interaction is lower than that of the αβ TCR (Mallis et al., 2015), suggesting the possibility that binding of the pre-TCR might require facilitation by other molecules. We have previously found that the interaction between stromal cells and developing T cells at the β-selection checkpoint involves recruitment of the microtubule organizing center (MTOC), a hallmark of initiation of the immunological synapse (Martín-Cófreces et al., 2008; Pham et al., 2015). This finding, combined with the interaction between pre-TCR and stromal cell MHC, opened the possibility that an immunological synapse might facilitate TCR signaling during β-selection.

Here, we demonstrate that the developing T cell indeed does form an immunological synapse upon interaction with stromal cells during β-selection. We characterize the synapse in *in vitro* T cell cultures and *in situ* in an intact mouse thymus. We further show that establishment of the synapse relies upon cooperation between pre-TCR and two key signaling pathways, which are essential for robust progression beyond the β-selection, Notch and CXCR4 signaling (Ciofani et al., 2004; Janas et al., 2010; Maillard et al., 2006; Trampont et al., 2010a; Wolfer et al., 2002). Finally, we demonstrate that the immunological synapse is a prerequisite for proliferation following the β-selection checkpoint.

## Results

### DN3 cells make an immunological synapse *in vitro* & *in situ*

Our previous observation that the MTOC was recruited to the interface with the stromal cell (Pham et al., 2015), reminiscent of an immunological synapse in mature cells (Alcover et al., 2016b; Dustin and Baldari, 2017), led us to assess whether developing T cells might also form an immunological synapse. Flow cytometry confirmed that pre-TCR components, including pre-TCR-alpha chain (pTα), TCRβ chain, LCK, LAT and PKCθ were expressed in DN3 cells. Phosphorylation of the TCR-associated kinase, LCK (pLCK^394^ and pLCK^505^ (active and inactive forms respectively) and ZAP70 (pZAP70) in DN3 cells indicated active signaling through the pre-TCR (Fig. S1). To assess the localization of pre-TCR components, we incubated *in vitro* generated DN3 cells with OP9-DL1 stromal cells, and analyzed conjugates by immunofluorescence. Blinded scoring indicated that pTα and TCRβ were polarized in most conjugates (61.4% and 84.6% respectively) (Fig. 1A & B). Engagement of the TCR receptor in mature T cells initiates downstream signal transduction with polarization of its downstream signaling molecules and phosphorylation of the immunoreceptor tyrosine-based motifs (Alarcon et al., 2011; Alcover et al., 2016b; Gaud et al., 2018). Similarly for the developing T cells, the majority of conjugates showed polarization of total LCK (86.5%), pLCK^394^ (71.6%), pLCK^505^ (86.8%), pZAP70 (52.1%) and LAT (75.6%), but not PKCθ (28.3%). Together, these findings indicate that developing T cells cluster T cell receptor components at the interface with the stromal cells during the TCRβ checkpoint.

**Figure 1.**
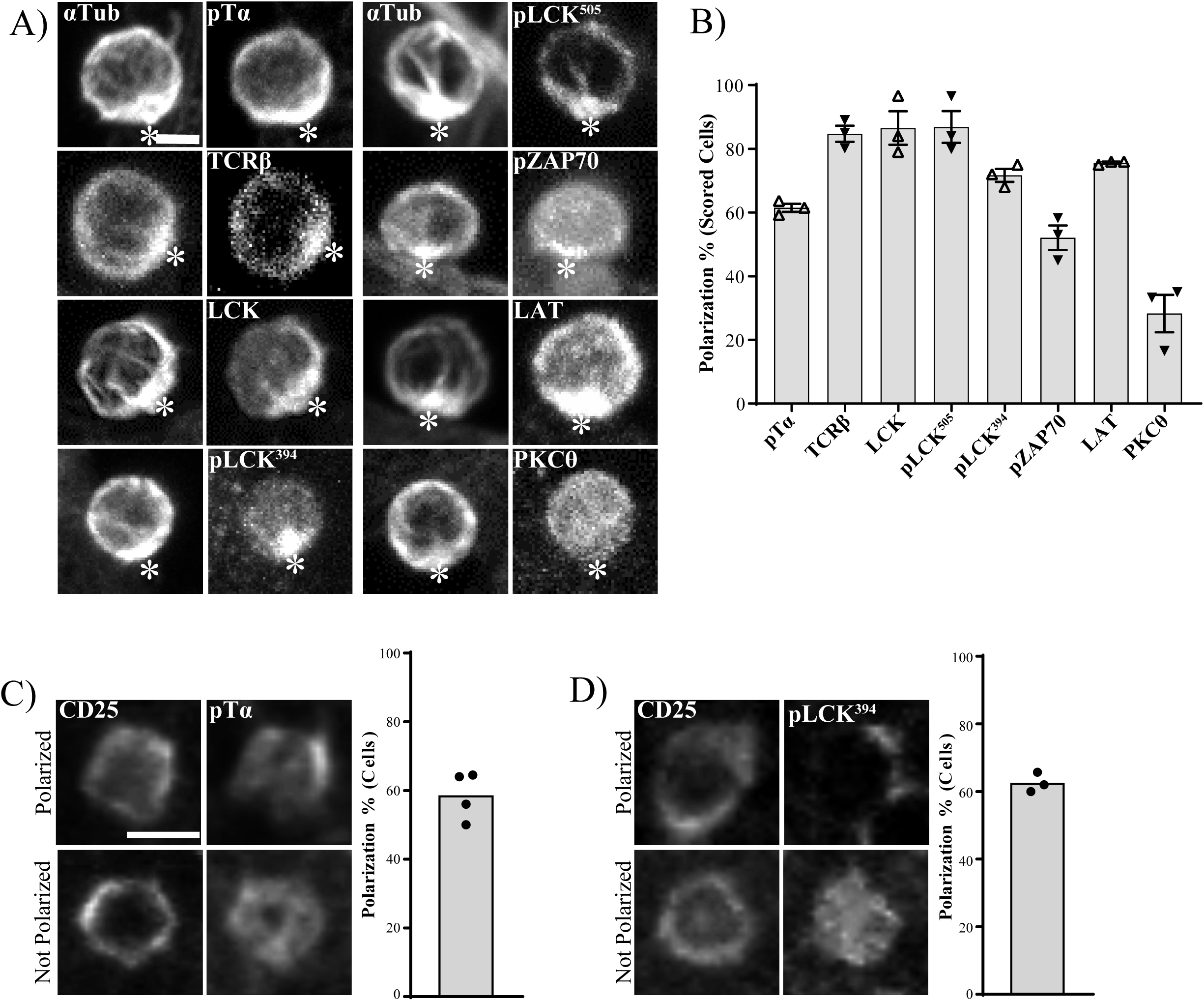
Polarization of pre-TCR components in DN3 cells *in vitro* and *in situ*. **A, B)** DN3 cells were incubated with OP9-DL1 cells for 14 hours, fixed and stained for α-tubulin to mark the MTOC, and a TCR-associated protein (as shown). Z-stack images were acquired using confocal microscopy, and representative images shown as maximum projections in (A). After triaging for cells in which MTOC was recruited to the interface with an OP9-DL1 cell, the percentage of cells in which the TCR component was polarized to the interface with the OP9-DL1 cell was determined by blind scoring (B). Total number of scored conjugates per marker is 75, 25 cells per biological replicate. Scale bar, 5µm. **C) & D**) DN3 cells in a section of an intact thymus were stained for CD25 as a non-polarized control, and either pTα (C) or pLCK^394^. Images were acquired using widefield fluorescent microscopy (Vectra® 3 automated quantitative pathology imaging system) and representative images of polarized (top row) and non-polarized (bottom row) are shown. Total number of scored cells n=156 (pTα) & n=134 (pLCK^394^). Scale bar, 5µm (A) and 10µm (C, D).

The above findings suggest that developing T cells form the equivalent of an immunological synapse. To ensure that these findings were not an artefact of the OP9-Dl1 stromal system, we assessed the localization of TCR components in an intact thymus. We used 6-color immunofluorescence imaging to investigate polarization of pTα, and pLCK^394^ in the DN3 population (Fig. 1C & D). pTα was polarized in 58.6%, and pLCK^394^ in 62.6% of DN3 cells in the thymus, while, as previously found (Pham et al., 2015) the cytokine receptor, CD25, was not polarized. These data suggest that developing T cells form an immunological synapse similar to that of mature T cells. This immunological synapse involves clustered pre-TCR and associated signaling molecules, and is present in T cells undergoing β-selection both *in vitro* and *in situ*.

### Notch1 and CXCR4 are required for immunological synapse formation *in vitro* and *in situ*

Given that the pre-TCR has only limited affinity for MHC-peptide complexes, we speculated that the recruitment of pre-TCR components to the interface with the stromal cell might depend upon other receptors. The fate determinant, Notch1, and the chemokine receptor, CXCR4 play essential roles in driving T cell development at the β-selection checkpoint (Maillard et al., 2006; Trampont et al., 2010a; Trampont et al., 2010b), and are localized at the DN3-OPL-DL1 stromal cell interface (Pham et al., 2015) (Fig. S2). Staining of DN3 cells conjugated to OP9-DL1 showed that pTα co-polarized with Notch1 (68.4%) and CXCR4 (80%) (Fig. 2A & B). We investigated whether this co-polarization could be observed in an intact mouse thymus using five color imunofluorescence imaging. Almost all DN3 cells in the intact thymus were polarized for Notch1 and CXCR4, so we selected DN3 cells that were clustered for pTα, and scored for whether the pTα cluster co-localized with Notch1 and CXCR4. pTα was co-clustered with Notch1 (65.3%) of these cells and with CXCR4 (59.5%) (Fig 2C & D). These data indicate that Notch1 and CXCR4 assemble with the TCR signaling platform, and raise the possibility that they facilitate the establishment of the immunological synapse.

**Figure 2.**
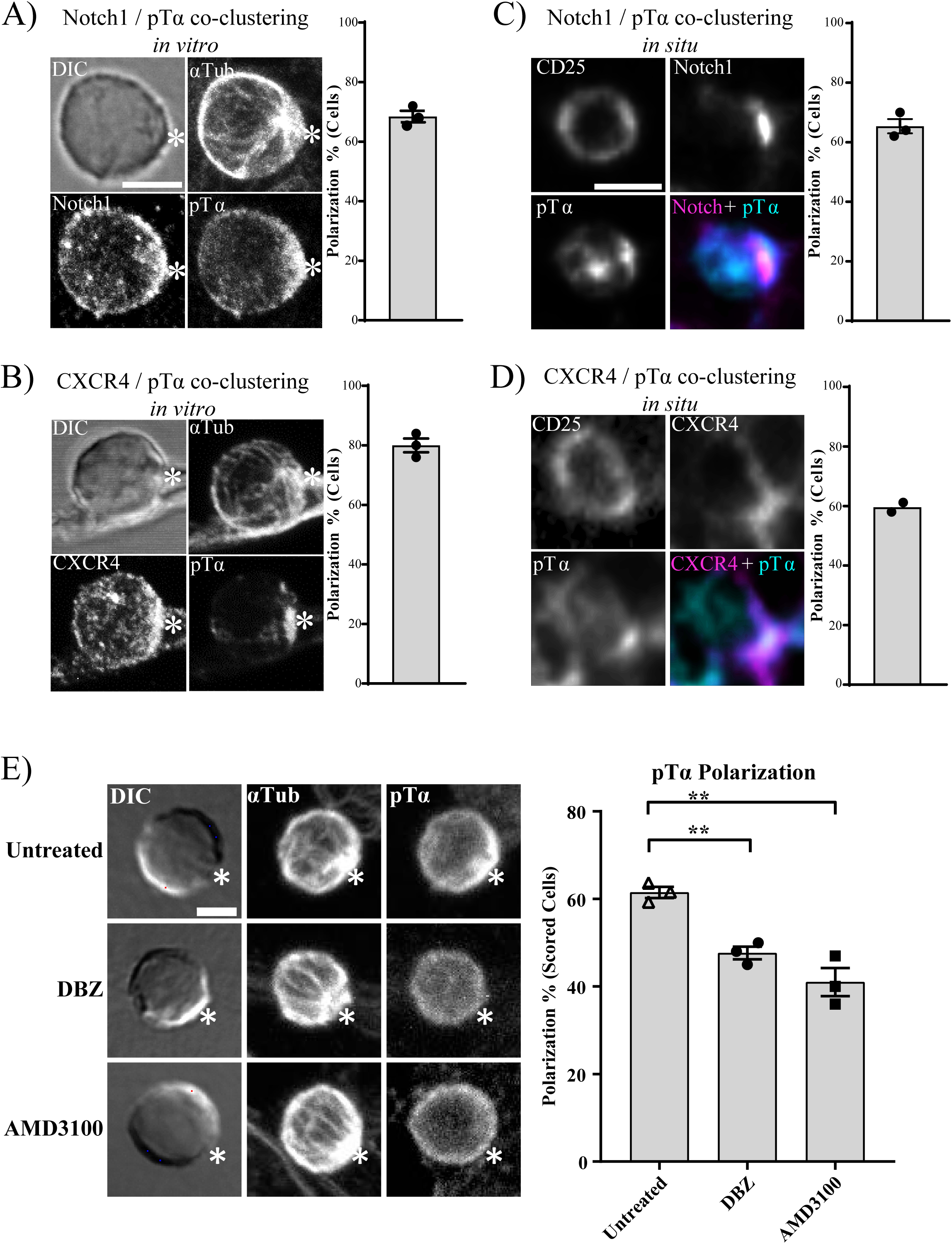
Notch and CXCR4 recruit pre-TCR components to the stromal interface. **A, B)** DN3 cells were incubated with OP9-DL1 cells for 14 hours, fixed and stained for α-tubulin to mark the MTOC, Notch1, CXCR4 and pTα chain (as shown). Z-stack images were acquired using confocal microscopy, and representative images shown as maximum projection in (A, B). White asterisk in A) & B) indicates the interface between DN3 cells and OP9-DL1 cells. After triaging for cells in which MTOC was recruited to the interface with an OP9-DL1 cell, the percentage of cells in which pTα chain was polarized with either Notch1 or CXCR4 to the interface with the OP9-DL1 cell was determined by blind scoring. Total number of scored conjugates per marker is 75, 25 cells per biological replicate. Scale bar, 5µm. **C) & D)** DN3 cells in an intact thymus were stained for CD25 as a non-polarized control, pTα as a marker for pre-TCR, and either Notch1 (C) or CXCR4 (D). Images were acquired using widefield fluorescent microscopy (Vectra® 3 automated quantitative pathology imaging system) and representative images of Notch1-pTα (C) and CXCR4-pTα (D) co-clustering *in situ* are shown. Total number of scored cells 150 (C) and 100 (D). **E)** DN3 cells were incubated with OP9-DL1 cells for 14 hours in the presence or absence of either Notch inhibitor (DBZ) or CXCR4 inhibitor (AMD3100), then fixed and stained for α -tubulin to mark the MTOC and pTα chain as a marker of pre-TCR (as shown). After triaging for cells in which MTOC was recruited to the interface with an OP9-DL1 cell, the percentage of cells in which pTα was polarized to the interface with the OP9-DL1 cell was determined by blind scoring (as shown by column-bar plot). Total number of scored conjugates per condition is 75, 25 cells per biological replicate. Scale Bar, 5µm (A & E), 10µm (C). Statistics: One Way ANOVA Test: Untreated vs DBZ, **P=0.019; Untreated vs AMD3100, **P=0.001.

The co-clustering of Notch1 and CXCR4 with pTα led us to suspect that Notch1 and CXCR4 might regulate pTα polarization. Indeed, scored DN3 conjugates treated for 14 hrs with pharmacological inhibitors of Notch signaling (DBZ) or CXCR4 signaling (AMD3100), showed 47% and 41% pTα chain polarization, respectively, compared to 63.6% in the untreated sample (Fig. 2E). We saw a similar reduction in pTα polarization after 3 hours of signaling inhibition, suggesting that this effect is direct, rather than an indirect consequence of phenotypic changes induced by Notch or CXCR4 signaling (Fig. S3). These data suggest that Notch1 and CXCR4 act as recruiters of the pTα chain to the interface as a part of the *de novo* assembly of the immunological synapse.

### The requirement for Notch1 and CXCR4 in passing the β-selection checkpoint is circumvented by replacing the immunological synapse with pharmacological mimicry of TCR signaling

Both Notch and CXCR4 cooperate with pre-TCR signaling to facilitate progression beyond the β-selection checkpoint (Chann and Russell, 2019), but the mechanisms by which these signaling pathways interact is not known. We assessed whether the influence of Notch1 and CXCR4 signaling on β-selection might at least in part, reflect their influence on the formation of the immunological synapse. It is well established that the β-selection checkpoint requires Notch1 and CXCR4 signaling (Ciofani et al., 2004; Janas et al., 2010; Maillard et al., 2006; Trampont et al., 2010a; Yashiro-Ohtani et al., 2009), and we confirmed the impact of inhibiting Notch and CXCR4 signaling in the OP9-DL1 co-cultures, where the survival of DN3a cells co-cultured with the Notch inhibitor, DBZ, or the CXCR4 inhibitor, AMD3100, was not affected, but fewer cells differentiated to DN3b and DP compared with the untreated co-culture (Fig S4A). We assessed whether the impact of these inhibitors on differentiation correlates with an immediate downstream marker of pre-TCR signaling, pLCK^394^. 3 hours after addition of Notch and CXCR4 inhibitors, pre-TCR signaling was reduced as indicated by pLCK^394^ levels (Fig S4B). The effectiveness of each inhibitor was confirmed by the reduction in expression of Hes1 and CXCR4 (readouts of Notch and CXCR4 signaling, respectively (Azab et al., 2009; Kageyama et al., 2007; Lee et al., 2013)). Thus, the long-term impact on differentiation correlated with a short-term impact on pre-TCR signaling.

The above results demonstrate a functional interaction between Notch1, CXCR4 and proximal pre-TCR signaling, but does not indicate whether that interaction occurs via assembly of the immunological synapse. To assess this, we turned to a pharmacological mimic of TCR signaling, phorbol 12-myristate 13-acetate (PMA), which bypasses the need for a signaling platform via directly activating PKCθ (Felli et al., 2004; Tahara et al., 2009). However, to avoid confounding the results with cells that had already assembled an immunological synapse, we sorted DN3a cells for the presence or absence of surface TCRβ. As previously published, the cell surface expression of TCRβ dramatically alters subsequent fate of the cell populations (Klein et al., 2019). After 5 days of co-culturing on OP9-DL1, TCRβ^+^ cells yielded more than double the cells yielded by TCRβ^-^ cells, with commensurate differences in CFSE dilution, indicating higher cell division rates (Fig. 3A). TCRβ^+^ cells produced increased numbers of DP cells compared with TCRβ^-^ cells, and fewer DN3a cells remained in the culture (Fig. 3B & C). In addition, Annexin V and PI staining showed that TCRβ^+^ DN3a yielded fewer apoptotic and necrotic cells (Fig. S5). Thus, the presence and assembly of the pre-TCR is associated with increased differentiation and proliferation, and TCRβ^-^ cells should represent TCR-signaling naïve cells appropriate for comparing canonical pre-TCR signaling with PMA-induced signaling.

**Figure 3.**
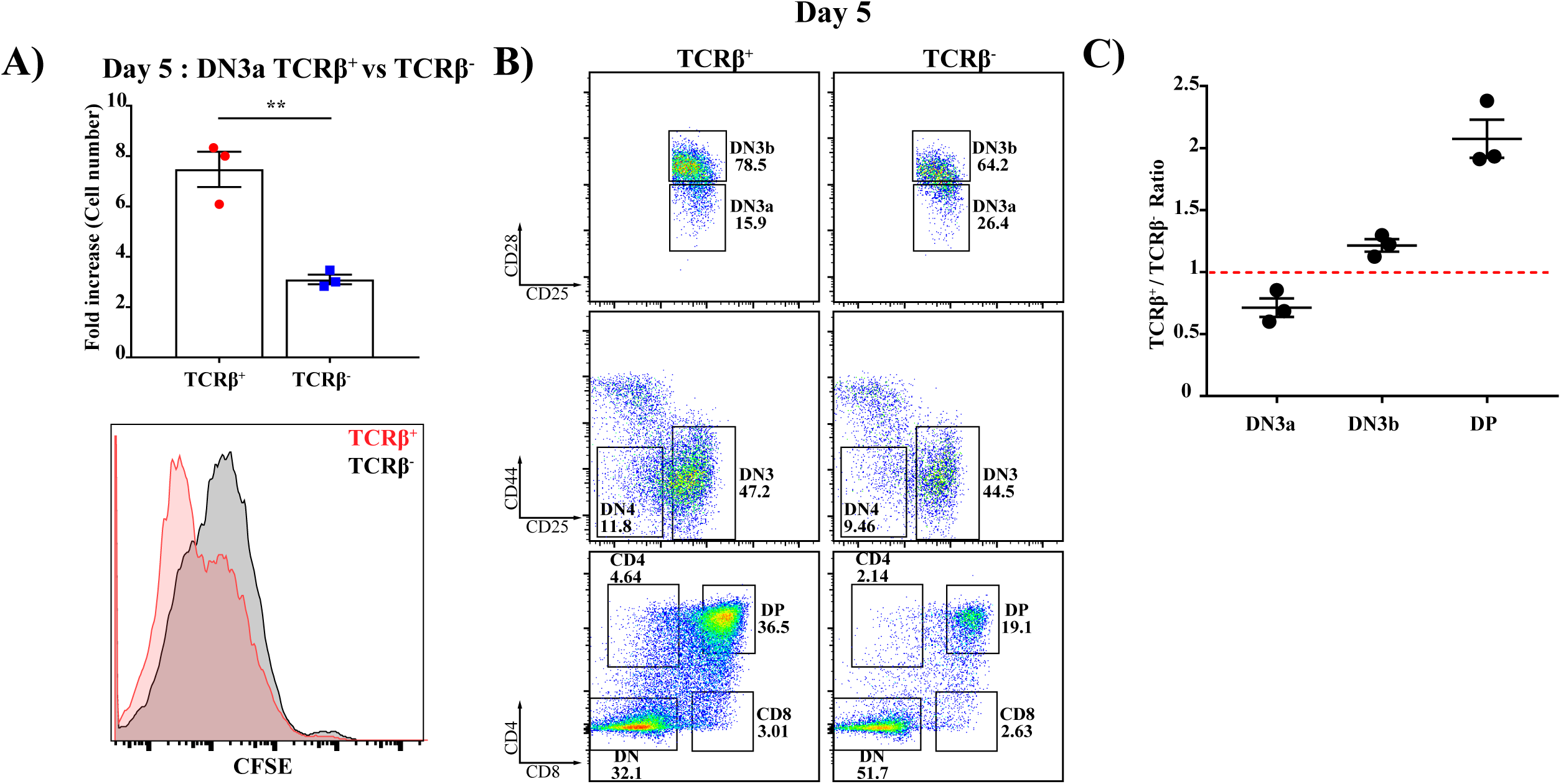
Pre-TCR assembly at DN3a is critical for transition beyond the β-selection checkpoint. Purified TCRβ^+^ and TCRβ^-^ DN3a cells were co-cultured with OP9-DL1 for 5 days. Cellularity was assessed by measuring fold increase in both co-cultures (as shown in the column-bar plot), and proliferation was assessed using the intensity of CFSE (as shown in histogram, bottom left). Flow cytometry was used to assess the differences between TCRβ+ and TCRβ-DN3a co-cultures progression and differentiation to subsequent developing stages (as shown by flow cytometry plots). The percentages of DN3a, DN3b and DP cells from 3 independent biological replicates were expressed as a ratio (TCRβ^+^/ TCRβ^-^) as shown by dot-plot (top right). Data presented are representative of 3 independent experiments. Statistics: Unpaired T-test, **P= 0.004.

To generate an initial population with no pre-assembled immunological synapse, we therefore sorted for TCRβ^-^ DN3a cells to explore functional interactions between TCRβ, Notch and CXCR4 signaling. A time course of response to incubation on OP9-DL1 cells revealed that TCRβ^-^ DN3a cells remain viable and proliferate over 48 hours (data not shown), but show very little progression through the differentiation stages, with most of the cells remaining as DN3a, suggestive of multiple cycles of self-renewal (Fig. 4I)). As expected, the addition of PMA caused differentiation through DN3b and DN4 (Fig. 4II). Remarkably, Notch inhibition had no or a small inhibitory effect on differentiation from DN3a without PMA treatment, but substantially increased differentiation from DN3a to DN3b and DN4 in the PMA treated cells (compare Fig. 4III with IV). The effects of Notch inhibition on differentiation were not related to cell death (Fig. S6). Inhibition of CXCR4 also showed a differential effect on differentiation in untreated versus PMA-treated cells, albeit to a lesser extent than inhibition of Notch (Fig. 4V & VI). These results combined indicate that, although both Notch and CXCR4 are required for differentiation in response to pre-TCR signaling, they are not required for differentiation in response to PMA signaling. Combined with our previous finding that Notch1 and CXCR4 are required for synapse formation and proximal TCR signaling, these results support the notion that, rather than directly impacting upon differentiation, the major role for Notch1 and CXCR4 during β-selection is in *de novo* assembly of the immunological synapse.

**Figure 4.**
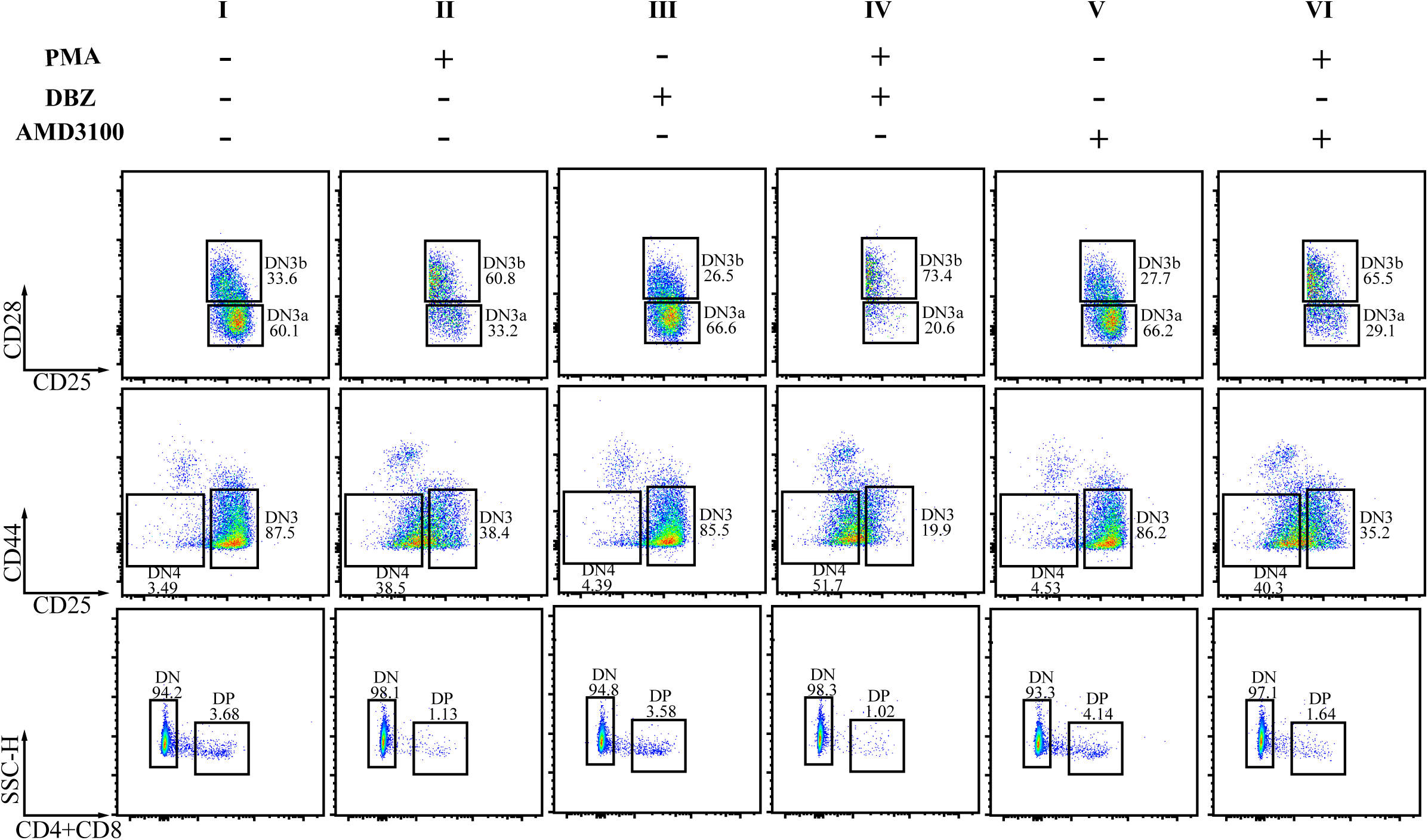
The effect of inhibition of Notch and CXCR4 signaling on progression beyond β-selection indicates a role in immunological synapse assembly. . Purified TCRβ^-^ DN3a cells were co-cultured on OP9-DL1 cells in the presence or absence of either of the following drug combinations; Notch inhibitor (DBZ), CXCR4 inhibitor (AMD3100), a pharmacological mimic of pre-TCR downstream signaling (PMA), PMA with Notch inhibitor and PMA with CXCR4 inhibitor. Co-cultures were stopped after 48 hours and analysed by flow cytometryto assess progression and differentiation (as shown by flow cytometry dot-plots). The presented data are representative of three independent biological replicates.

### The co-assembly of Notch1, CXCR4 and pre-TCR into a single signaling platform is correlated with progression through the β-selection checkpoint *in situ*

It is well established that signaling through the pre-TCR during β-selection promotes cell proliferation (Miyazaki et al., 2008; Petrie et al., 2000). Therefore, we hypothesized that formation of the immunological synapse could be required for cell proliferation after β-selection. Hence, we investigated the correlation between the co-clustering of Notch1 and CXCR4 with pLCK^394^ and proliferation of DN3 cells *in situ* (Fig. 5). We used 6-colour immunofluorescent staining of thymus sections, with Ki67 as a proliferation marker. Notch1 was co-clustered with pLCK^394^ in 63.3% of the scored cells in the subcapsule. Interestingly, 70.9% of the cells that showed co-polarization of Notch1 and pLCK^394^ at subcapsule expressed Ki67, compared to 16.2% of these that did not show co-polarization (Fig. 5A & C). Similarly, CXCR4 was co-clustered with pLCK^394^ in 68% of the scored cells in the subcapsule. As with Notch1, 80.4% of the cells that showed co-polarization of CXCR4 and pLCK^394^ at subcapsule expressed Ki67, compared to 37.4% of these that did not show co-polarization (Fig. 5B & D). To be certain that Notch1 and CXCR4 co-polarization with pLCK^394^ is essential for proliferation and not due to pLCK^394^ clustering, we scored cells that showed clustering of pLCK^394^ only. Strikingly, pLCK^394^ clustering did not correlate with Ki67 expression in the subcapsule (Fig. 5E). These data strongly support the hypothesis that Notch1 and CXCR4 are required for a proper assembly of the pre-TCR immunological synapse, which in turn, propagates signaling via pLCK^394^ to promote proliferation. These data together indicate that proliferation at the β-selection checkpoint is associated with formation of a signaling platform at the stromal interface, comprising Notch1, CXCR4, and the pre-TCR complex.

**Figure 5.**
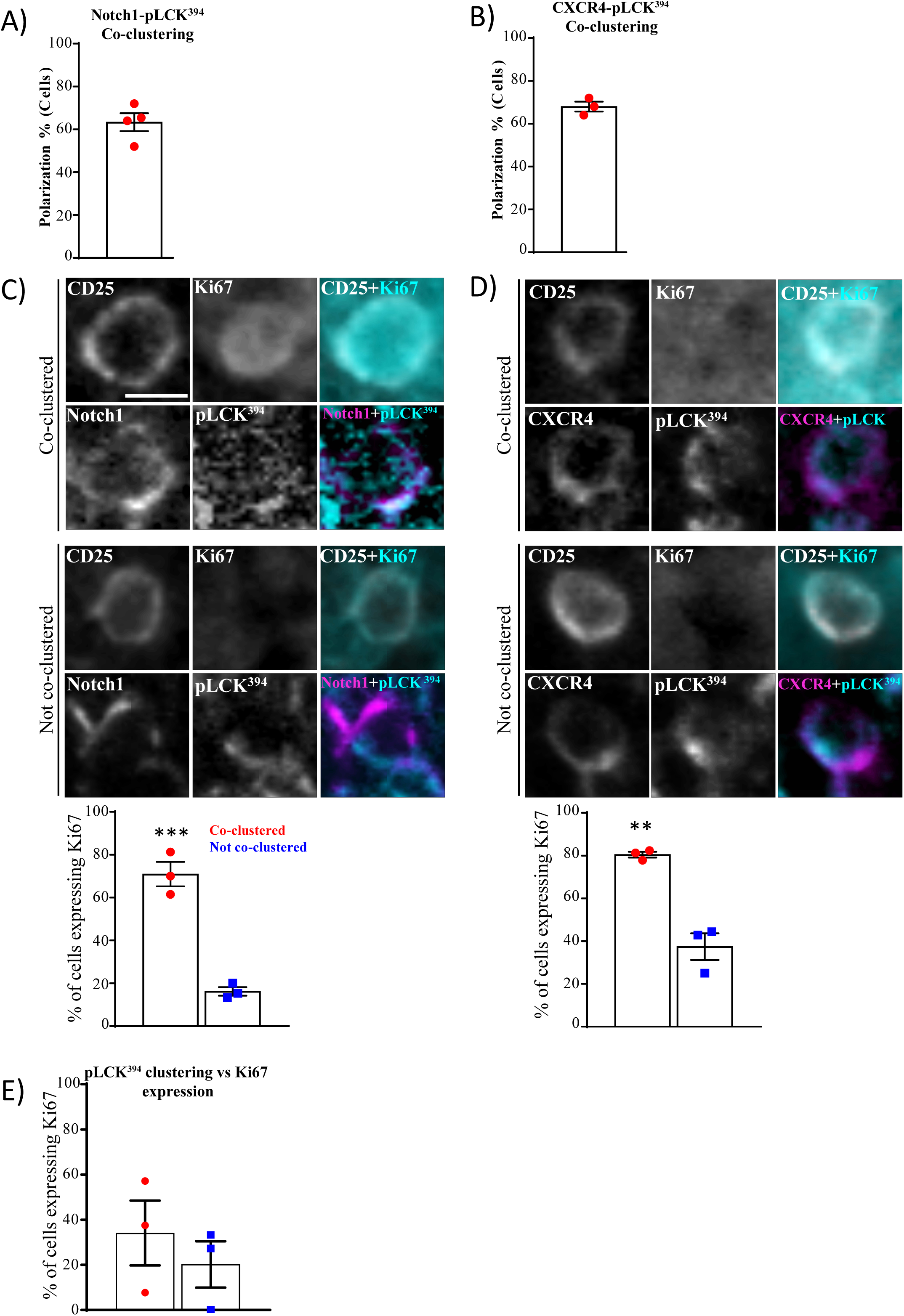
The co-assembly of Notch1, CXCR4 and pre-TCR into a single signaling platform is strongly associated with proliferation at the subcapsule region *in situ*. A, B) Multiplex imaging on sections of an intact thymus was carried out, followed by automated tissue classification using HALO software. DN3 cells in the subcapsule ere identified, and analysed for CD25 as a non-polarized control. The percentages of cells showed co-clustering of pLCK^394^ as a marker of pre-TCR signaling with either Notch1 (A) or CXCR4 (B) was determined as shown by the column-bar plots. Total number of scored cells is 100 (A) and 75 (B), with 25 cells per biological replicate. **C, D)** DN3 cells were stained for CD25 as non-polarizing control, pLCK^394^ as marker for pre-TCR downstream signaling, Ki67 as a proliferation marker and either Notch1 or CXCR4. Images were acquired using widefield fluorescent microscopy (Vectra® 3 automated quantitative pathology imaging system) and representative images of highly expressed Ki67 when Notch1 (C) or CXCR4 (D) co-cluster with pLCK^394^ or lack of Ki67 expression when Notch1 or CXCR4 do not co-cluster with pLCK^394^ *in situ* are shown. DN3 cell with co-clustered or no co-clustering of pLCK^394^ with either Notch1 (C) or CXCR4 (D) in the subcapsule were assessed for the levels of Ki67 expression as shown by column-bar plots. Total number of scored cell is 75, with 25 cells per biological replicate.. **E)** DN3 cells in an intact thymic section were identified in the subcapsule and the percentages of cells expressing the proliferation marker, Ki67, were assessed in cells showed clustering or no clustering of pLCK^394^ as shown by column-bar. Total number of scored cells is 75, with 25 cell per biological replicate. Scale bar, 10µm. Statistics: Unpaired T-Test, **P=0.0025, ***P=0.0008.

### MHC plays a role in pTα chain clustering and the establishment of the pre-TCR immunological synapse

In mature T cells, the establishment of the immunological synapse is ligand-dependent (Dustin et al., 2010). Although the pre-TCR signaling is ligand-independent, it was shown recently that the pre-TCR can interact with peptide-MHC complexes to influence T cell development (Mallis et al., 2015). Therefore, we assessed whether a physical association of MHC and pre-TCR took place in an intact thymus. Using 6-color immunofluorescent staining, we first assessed the proximity of DN3 cells to MHC Class II (which we refer to as MHC)-expressing cells in intact thymic lobes. We randomly picked five different regions of the same size the in thymic lobe (two thymic lobes from different mice, with a total of 322 scored cells) and scored cells for being either adjacent to or distant from MHC. We found that 52% of DN3 cells were adjacent to MHC (Fig. 6A). pTα was clustered in a higher proportion of DN3 cells adjacent to MHC (87.8%) as compared to DN3 cells distant from MHC (61.4%) (Fig. 6B). Moreover, of the DN3 cells adjacent to MHC and with clustered pTα, the clustered pTα was oriented towards MHC in 82.7% (Fig. 6B). These data suggest that the pre-TCR can interact with MHC in the intact thymus, supporting previous published work, suggesting that interactions with peptide-bound MHC fosters expansion of cells that can recognize self-MHC (Mallis et al., 2015). Moreover, these results suggest that MHC interactions are associated with the establishment of an immunological synapse during β-selection.

**Figure 6.**
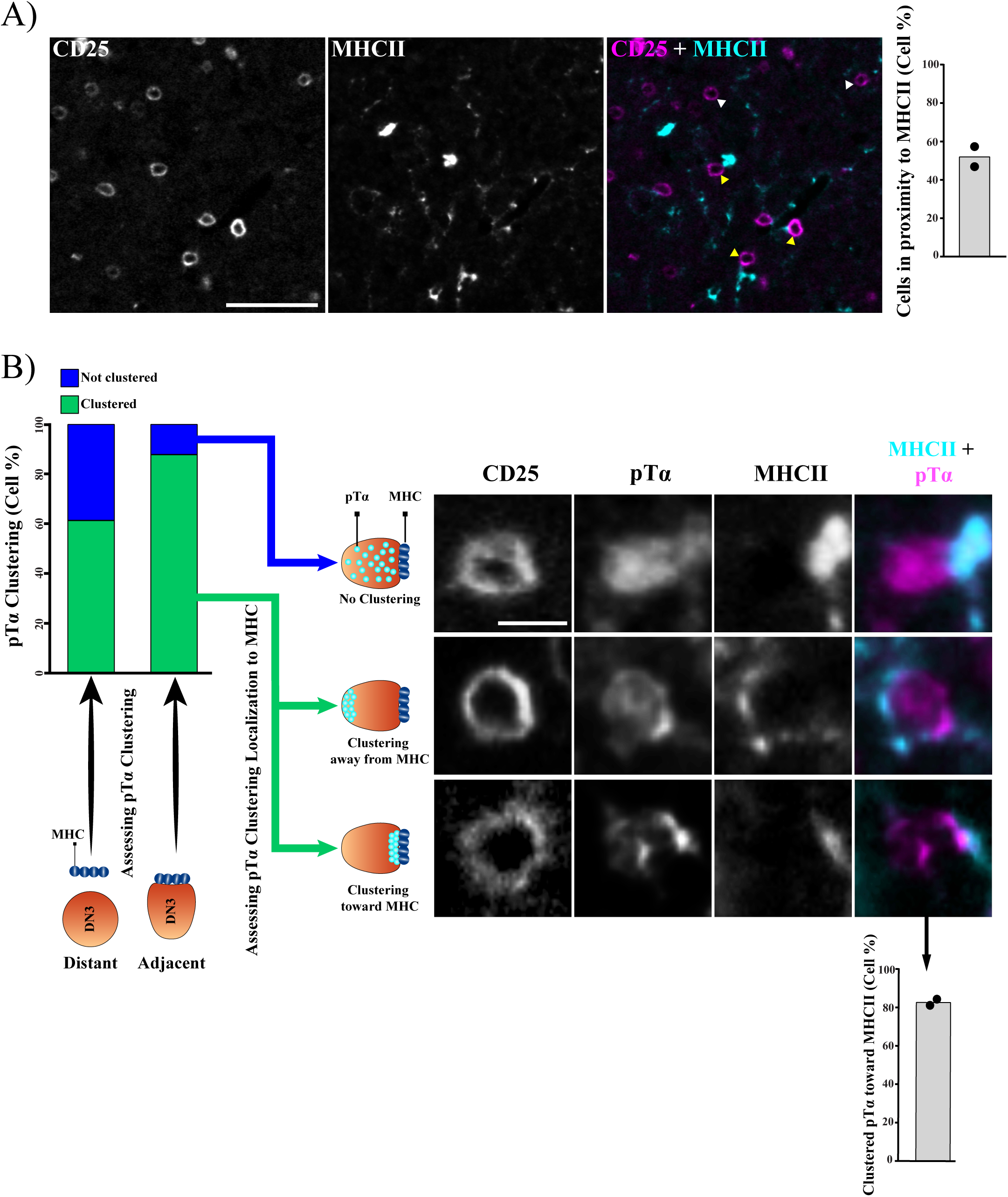
MHC plays a role in establishing the pre-TCR immunological synapse at DN3 stage *in situ.* **A)** DN3 cells were identified in 5 randomly picked regions of the same size in an intact thymus to assess their localization to MHCII. Images were acquired using widefield fluorescent microscopy (Vectra® 3 automated quantitative pathology imaging system) and are representative of DN3 cells adjacent (yellow arrow heads) or away (white arrow heads) from MHCII. DN3 cells were then scored as either adjacent to MHCII or away from it, as shown by the column-bar plot.. Total number of scored cells 322, with 161 per biological replicate. **B)** pTα clustering was assessed in DN3 cells adjacent to or away from MHCII as shown in the stacked column-bar plot. DN3 cells adjacent to MHCII, and that showed pTα clustering, were assessed for the localization of pTα clustering as compared to MHCII as shown by column-bar plot. Images were acquired using widefield fluorescent microscopy (Vectra® 3 automated quantitative pathology imaging system) and representative image of DN3 cells adjacent to MHCII with clustering of pTα toward MHCII (I), clustering of pTα away from MHCII (II) and no clustering of pTα (III) are shown. Scale bar, 50µm (A); 10µm (B).

## Discussion

The β-selection checkpoint is the stage at which the developing T cell tests whether it has effectively recombined the gene for TCRβ (Bednarski and Sleckman, 2012; Rothenberg et al., 2008). Cells that fail this test will die, and cells that can signal through the pre-TCR downstream will pass the test and survive, proliferate and differentiate (Kreslavsky et al., 2012; Michie and Zuniga-Pflucker, 2002). In this study, we show that this test of pre-TCR signaling involves the establishment of an immunological synapse, resembling the immunological synapse of mature T cells (Fig. 7). The immunological synapse provides a platform for downstream signaling and progression beyond the β-selection checkpoint. Differing from that of mature T cells, the β-selection immunological synapse is facilitated by Notch1 and CXCR4 signaling.

**Figure 7.**
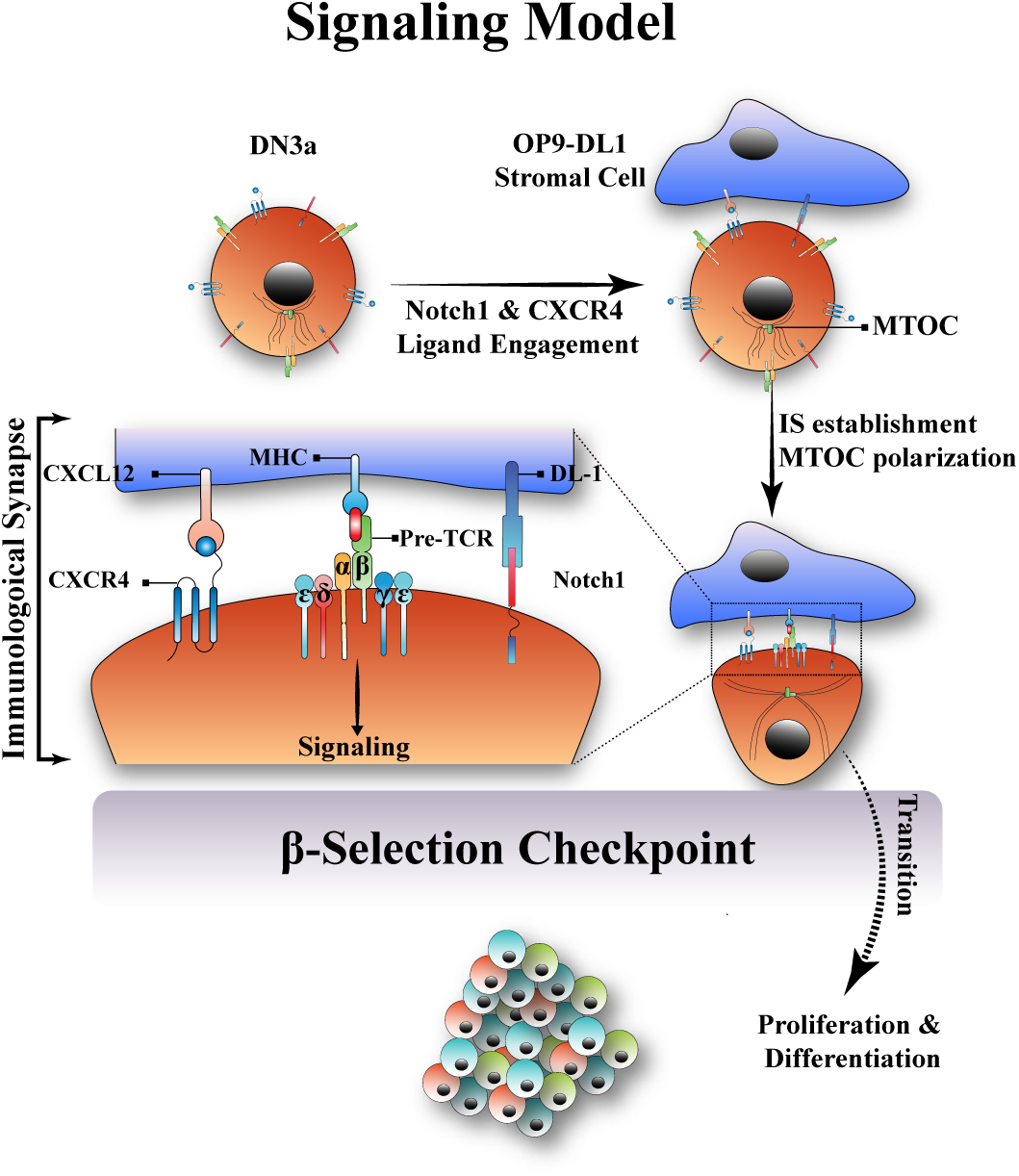
A new model for the β-selection checkpoint that incorporates an immunological synapse. In the proposed model, contact of DN3a) developing T cells with stromal cells in the thymus (OP9-DL1 in ***in vitro)*** causes Notch1 and CXCR4 to engage with their ligands (DL1/4 and CXCRL12, respectively). Notch1 and CXCR4 engagement promotes the establishment of the immunological synapse via the pre-TCR (pTα and TCR-β chains). The immunological synapse is correlated with the polarization of MTOC, clustering of the CD3 complex clustering (ε, δ and γ chains) and proximity to MHC on the stromal cell. The immunological synapse then provides the signaling platform required for downstream signal transduction through the pre-TCR. Signaling via the pre-TCR promotes transition beyond the β-selection checkpoint, followed by proliferation and differentiation to the subsequent T cell developmental stages.

The immunological synapse in a mature T cell is characterized by the polarization of the TCR components toward the antigen presenting cell (APC) (Dustin et al., 2010; Martín-Cófreces et al., 2008; Mittelbrunn et al., 2011). In a similar manner, we find that the MTOC polarizes at the interface between DN3 and OP9-DL1 stromal cells (Pham et al., 2015), along with pre-TCR structural and signaling components such as Lck and LAT. In addition, using OPAL multiplex imaging *in situ*, we also show polarization *in situ* in the intact thymus, of both pTα (a component of the pre-TCR complex) and pLCK^394^ (an indicator of signaling downstream of the pre-TCR). These results indicate that a signaling platform akin to the well characterized immunological synapse seen in mature T cells occurs during β-selection.

An intriguing aspect of the β-selection immunological synapse is the differences implied by incorporation of the pre-TCR, rather than the αβ TCR. In the mature T cell immunological synapse, signaling and polarization is driven by TCR engagement with peptide-bound MHC (Fooksman et al., 2010). The possibility of the pre-TCR binding to peptide-bound MHC has only recently been indicated based upon interactions with purified proteins (Mallis et al., 2015). In support of a physiological relevance for binding to MHC, the pre-TCR-MHC interaction was correlated with elevated levels of calcium influx, indicating active signaling (Das et al., 2016; Mallis et al., 2015). This interaction is clearly not essential for T cell development (Irving et al., 1998; Koller et al., 2010), but by encouraging the preferential proliferation of cells whose pre-TCR can bind self-MHC, is thought to skew the repertoire towards TCRβ-repertoires before undergoing TCRα rearrangement (Mallis et al., 2015). Our findings that MHC and pre-TCR co-cluster towards each other suggest that the binding has functional relevance in the intact thymus. The formation of an immunological synapse also provides a potential mechanism for overcoming the weak binding between the pre-TCR and peptide-bound MHC, particularly before expression of the co-receptors, CD4 and CD8. Together, these observations suggest that, although the β-selection immunological synapse does not depend upon peptide-bound MHC for its assembly, the synapse might promote pre-TCR signaling in response to peptide presented by adjacent thymic stromal cells.

Notch1 and CXCR4 are included in the β-selection immunological synapse *in vitro* and *in situ*, and their signaling is required for optimal assembly. These findings are compatible with our previous observations that Notch1 and CXCR4 polarize at the interface between DN3 cell and OPL-DL1 (Pham et al., 2015). Others have identified a functional interaction between the pre-TCR and either Notch (Ciofani et al., 2004; Yashiro-Ohtani et al., 2009) or CXCR4 (Trampont et al., 2010a), and that Notch and TCR are co-recruited to the immunological synapse of mature T cells and DP (CD4^+^CD8^+^) thymocytes (Anderson et al., 2005; Guy et al., 2013). In mature T cells, TCR signaling enhances Notch recruitment to the immunological synapse (Guy et al., 2013). In contrast, our findings suggest that Notch and CXCR4 signaling are required for optimal assembly of a pre-TCR-containing immunological synapse. It is possible that the requirement for Notch and CXCR4 reflects compensation for the weaker binding of the pre-TCR to MHC, by creating a stable interface between the cells that allows for formation of the immunological synapse. Supporting the possibility that Notch and CXCR4 signaling initiate polarization, the MTOC (a key component of the immunological synapse) is recruited to the interface in response to both Notch and CXCR4 signaling.

These findings point to a possible alternative explanation for the cooperation between Notch, CXCR4 and pre-TCR at the β-selection checkpoint: that in addition to (or instead of) cooperating at the level of downstream signaling, Notch and CXCR4 act to promote pre-TCR signaling by enabling the immunological synapse. To test this theory, we took advantage of a well characterized surrogate for TCR signaling that does not involve assembly of an immunological synapse, PMA (Felli et al., 2004; Tahara et al., 2009). To compare PMA treatment with pre-TCR signaling, we established a system in which cells either preferentially progressed though the β-selection checkpoint via PMA signaling (DN3a cells that had not yet expressed the pre-TCR) or only via signaling through the pre-TCR (a mixed population of DN3a cells both expressing and not expressing TCRβ that were not treated with PMA). Consistent with recently published work (Klein et al., 2019), DN3a cells that expressed cell surface TCRβ differentiated faster and proliferated more than those that did not express TCRβ, compatible with the pre-TCR as being a prerequisite for progression beyond the β-selection checkpoint. We used this system to discriminate between a role for Notch and CXCR4 in immunological synapse assembly compared with downstream signaling. The diametrically opposed responses of the two forms of pre-TCR signaling to inhibition of either Notch1 or CXCR4 suggest that, at β-selection, Notch1 and CXCR4 effects on pre-TCR signaling act via the immunological synapse.

A hallmark of pre-TCR signaling is the onset of proliferation (Aifantis et al., 2001; von Boehmer, 2005). Accordingly, we tested if the co-clustering of Notch1 and CXCR4 with the pre-TCR downstream active signaling marker, pLCK^394^ correlated with proliferation of DN3 cells. Remarkably, our *in situ* multiplex imaging showed that co-clustering of Notch1 and CXCR4 with pLCK^394^ in DN3 thymocytes at the subscapular region had significantly higher expression of the proliferation marker, Ki67, compared to those that showed no co-clustering. This indicates that Notch1 and CXCR4 co-clustering with pLCK^394^ is critical for the proliferative expansion onset following pre-TCR signaling. Moreover, the pLCK^394^ clustering itself did not show strong correlation to proliferation, which further supports the essential role of Notch1 and CXCR4 in establishing this signaling platform. Together, these findings suggest that assembly of the pre-TCR into an immunological synapse is a precondition for proliferation in response to β-selection.

In conclusion, we demonstrate the establishment of an immunological synapse that facilitates pre-TCR signaling during the β-selection checkpoint. We show that this immunological synapse is regulated cooperatively by Notch1 and CXCR4 signaling. We propose that the β-selection immunological synapse is critical for regulation of pre-TCR signaling and promotes progression beyond the β-selection stage.

## Materials and Methods

### Primary hematopoietic co-culture

The OP9 stromal cell line transduced to express the Notch ligand, Delta-like1 (OP9-DL1) (Schmitt and Zuniga-Pflucker, 2002) was cultured in Minimal Essential Medium Alpha Modification (SAFC Biosciences, Sigma Aldrich) supplemented with 10% (v/v) Fetal calf serum, glutamine (1mM, GIBCO-BRL) and 100 ng/mL penicillin/streptomycin, at 37°C, 10% CO2. Fetal liver cells were extracted from E14.5 C57 black 6 (C57Bl/6) mouse and used as hematopoietic stem cell (HSC) progenitors. OP9-DL1 stromal cells were seeded in 6-well plates and mouse fetal liver cells were seeded onto them in a ratio 1:1 (2×10^5^ cell/well). The co-culture was maintained at 37°C, 10% CO2 in Minimal Essential Medium Alpha Modification (SAFC Biosciences, Sigma Aldrich) supplemented 10% fetal calf serum (v/v), glutamine (1mM), β-mercaptoethanol (50µM, Calbiochem), sodium pyruvate (1nM, GIBCO-BRL), HEPES (10mM, GIBCO-BRL), 100 ng/mL penicillin/streptomycin, 1ng/mL mouse interleukin 7 (Peprotech) and 5ng/mL mouse FMS-like tyrosine kinase 3 (Peprotech). Upon hematopoietic confluency every 5-8 days, the co-culture was harvested via pressure pipetting or gentle scrapping, and then lymphocytes were separated from OP9-DL1 via pulse spin at 1400 rpm, and seeded onto fresh OP9-DL1 stromal cells in 6-well plates. To investigate the effect of Notch1 and CXCR4 on the differentiation of purified DN3a, DN3a cells were purified by flow cytometric sorting, co-cultured on OP9-DL1 stromal cells, and differentiation monitored in the presence and absence of Notch1 inhibitor, DBZ (1nM), CXCR4 inhibitor AMD3100 (2ug/mL) and the pharmacological mimic of TCR signaling, 12-phorbol-13-myristate acetate, PMA (40ng) (Sigma-Aldrich).

### Flow cytometry sorting and analysis

All antibodies were purchased from eBiosciences or BD Pharmingen unless otherwise specified. DN1-4, DP and SP thymocytes were distinguished using surface receptors, where DN1 (CD25^-^ /CD44^+^/CD4^-^/CD8^-^), DN2 (CD25^+^ /CD44^hi^/CD4^-^/CD8^-^), DN3 (CD25^+^/CD44^lo^/CD4^-^/CD8^-^), DN4 (CD25^-^ /CD44^hi^/CD4^-^/CD8^-^), DP (CD4^+^/CD8^+^) and SP (CD4^+^/CD8^-^ or CD4^-^/CD8^+^). DN3a (early) and DN3b were discriminated using the surface expression of CD28. We further discriminated late DN3a from early DN3b using TCRβ surface expression. Viability of cells was analyzed using FITC-Annexin V and/or Propidium iodide (PI). Refer to Table 1 for all antibodies.

### Confocal Microscopy and Immunofluorescence staining

OPL-DL1 stromal cells were seeded in each well of an 8-chamber slide (Thermofisher) at 3 × 10^3^ cell per well, and incubated overnight at 37°C and 10% CO2. 4-5 × 10^4^ hematopoietic precursors were seeded upon the OP9-DL1 stromal cells in each well with fresh media, and then left in the incubator for 14 hrs. Cells were then fixed with 3.7% (w/v) paraformaldehyde in 100 mM PIPES, 5 mM MgSO4, 10 mM EGTA and 2 mM DTT (20 min, Room Temperature (RT)), washed twice, and then permeabilised in 0.1% Triton X-100 in PBS without MgCl2 or CaCl2 (7 min, RT), then washed twice with PBS. All cells were blocked with 2% Bovine Serum Albumin (30 mins) (Sigma-Aldrich), then washed twice with PBS, and primary antibody was added and cells were left for 1 hr at RT on rocker. Cell were washed twice with PBS and secondary antibodies were added and cells were left for 1hr at RT on rocker, then cells were washed twice with PBS and mounted using Prolong Gold antifade (Molecular Probes). The slides were examined at room temperature using a FluoView FV1000 BX61 confocal microscope (Olympus) using a 40x oil immersion objective. 9-12 Z-stack images were acquired of a distance 1µm per step, then maximum intensity projections were generated using Image-J software. Refer to Table S1 for all used antibodies.

### Formalin-fixed paraffin-embedded (FFPE) tissue samples and Immunohistochemistry (IHC)

The thymi of 6 week old (C57Bl/6) mice were extracted and washed in PBS, then fixed in 10% Formaldehyde buffer for 18 hrs at RT. Fixed tissues was washed times in PBS, then embedded by histologist in paraffin. A serial section of 4µm thickness was preformed and slices were mounted on glass slides. After deparaffinization, heat antigen retrieval method using pressure cooker was used, the slides were placed in a plastic container filled with antigen retrieval buffer, 1mM Ethylenediaminetetraacetic acid (EDTA) (Sigma-Aldrich) solution, pH 9.0. To validate the specificity and detection of primary antibodies, chromogenic IHC was carried out for each single antibody as follows; slides were blocked using 2% BSA for 1 hr at RT on a rocker. Primary antibodies were incubated for 1hr at RT on a rocker, followed by washing twice with 1xTris-Buffered Saline 0.5% Tween (TBST) (1mM Tris Base, 1.8% NaCl, 0.5% Tween 20, pH 7.4). Endogenous peroxidase was blocked by incubating the slides in 0.3% hydrogen peroxidase for 10 mins at RT, followed by washing twice in 1xTBST. Horseradish peroxidase (HRP) conjugated secondary antibodies (Vector Labs) was added for 1hr at RT on a rocker, then slides were washed twice with 1xTBST. Slides were incubated with 3,3’-Diaminobenzidine (DAB) chromogen kit (DAKO) for 10 mins at RT on a rocker, then washed twice with 1xTBST. Samples were counter-stained with hematoxylin and dehydrated with ethanol and xylene to prepare for mounting. Slides were scanned using 40X air objective on Olympus V120 slide scanner (Fig. S7). The antibodies were used to build a thymic multiplex antibodies panel (Table S2)

### OPAL multiplex immunofluorescence imaging of thymic sections

#### Uniplex Immunofluorescence Validation

Following chromogenic detection, each of the assessed antibodies was further titrated for the OPAL multiplex imaging. Mono-color staining of each of the antibodies was performed using the OPAL 7 color kit (PerkinElmer), using the same protocol as IHC, and after the incubation with the HRP-conjugated secondary antibodies, slides were incubated with individual tyramide signal amplification (TSA)-conjugated fluorophores for 5-10 mins at RT temperature on a rocker, then washed three times with 1xTBST. Slides were then mounted with Citifluor™ Poly (vinyl pyrrolidone) plus antifadent Solution (Citifluor), and scanned at RT using 20X objective on the Vectra® 3 automated quantitative pathology imaging system (PerkinElmer) (Fig. S8). Acquired images were used to build spectral library of all the target proteins using inForm Cell Analysis software (PerkinElmer).

#### Multiplex Imaging

Following validating target antibodies and building the spectral library described in the paragraph above. Multiplex staining was performed by following the same steps for uniplex staining, and after adding the first TSA dye, the slides were placed in plastic container and microwaved for 1 min at 100°C, then microwaved for 10 mins at 75°C, to remove the primary and secondary antibodies, and leave behind the TSA dye, then the same steps were performed for the following antibodies. After the last antibody, slides were incubated with DAPI (1:1000 in 1xTBST) stain (Spectral DAPI, PerkinElmer) for 2-3 mins at RT on a rocker, then washed twice with 1xTBST. Slides were then mounted using Citifluor™ Poly (vinyl pyrrolidone) plus antifadent Solution, and the entire tissue was imaged using 20X objective on the Vectra® 3 automated quantitative pathology imaging system.

#### InForm and HALO software

Acquired images were loaded to inForm cell analysis software and the spectral library built from uniplex staining of the target antibodies was used to spectrally dissect multiplex images. Images were then exported as component .tiff files. Component .tiff files were then uploaded to HALO image analysis software (Indica Labs) and fused to build the thymic lobe. Using the HALO Highplex FL region classifier module, the software was trained to distinguish three main regions in the thymus, medulla, cortex and subcapsular region by manually picking 5-10 regions for each, then running automated classification on the entire tissue to highlight these three regions (Fig. S9). DN3 cells were identified either in the cortex or subcapsule using CD25^+^ CD44^-^ CD4^-^ CD8^-^ surface expression.

### Clustering and co-localization quantification

#### In vitro scoring and cell triaging

One of the hallmarks of the immunological synapse formation in mature T cells is the polarization of MTOC to the interface (Martin-Cofreces et al., 2008). Accordingly, we first identified cell conjugates *in vitro* by relying on the polarization of MTOC to the interface with OP9-DL1 stromal cells. We took in consideration that MTOC and OP9-DL1 should be on the same focal plane and only selected conjugates with visible fluorescence for the marker of interest. It should be noted that almost all of the markers in this study showed clusters. We scored for single marker clustering, where clustering at the interface was given score of 1 and no clustering was given score of 0. For double marker scoring, we scored for conjugates showing visible fluorescence intensity of both markers, at the same focal plane. We scored for the co-localization of both markers clusters at the interface, where co-localized clusters at the interface were given score of 1 and no co-localization at the interface was given score of 0.

#### In situ scoring and cell triaging

Due to the technical difficulty of using MTOC polarization as guideline to identify conjugates in our *in situ* system, we only scored for clustering and co-localizations of clustered markers of interest. First, DN3 cells were identified either in the cortex or subcapsule regions using CD25^+^ CD44^-^ CD4^-^ CD8^-^ surface expression. For single marker clustering, we only scored cells with visible fluorescence intensity of marker of interest and in focus. These cells were scored as 1 when they were clearly clustered, and as cells that showed clustering were given a score of 1; and no clustering, a score of 0. When comparing the clustering of more than one marker,: we scored cells with visible fluorescence intensity of both markers and in focus. We took the following strategy to avoid confounding the results with co-localization of non-clustered protein: Since almost all cells *in situ,* showed clustering of Notch1 (100% of cells) and CXCR4 (80-88% of cells), but fewer cells showed clustering of pTα (53%) and pLCK^394^ (61%), we only scored cells that showed clustering of pTα and pLCK^394^. We then scored the clustered markers for co-localization, where co-localized clusters were scored as 1 and non co-localized clusters scored as 0

## Supplemental Material

Supplemental Material includes Supplemental Figures 1-9, and Tables 1 and 2, which describe antibodies used in the studies.

## Acknowledgements

We thank Helena Richardson (La Trobe University) and Joe Trapani (Peter MacCallum Cancer Centre) for comments on the manuscript. This work was performed in part at the Biointerface Engineering Hub @ Swinburne, part of the Victorian node of the Australian National Fabrication Facility (ANFF), a company established under the National Collaborative Research Infrastructure Strategy to provide nano-and micro-fabrication facilities for Australia’s researchers. The multiplexed immunohistochemistry was supported by the Centre for Advanced Histology and Microscopy at the Peter MacCallum Centre, and the flow cytometry was enabled by support form the L.E.W. Carty foundation This work was funded by support from the Australian Research Council (FT0990405 to SMR), the National Health and Medical Research Council (APP1099140 to SMR), the Schweizerischer Nationalfonds zur Förderung der Wissenschaftlichen Forschung (Swiss National Science Foundation, SNSF) (grants PA00P3_142120 and P300P3_154664 to MC), and a Swinburne University Postgraduate Research Award to AA.

## Author Contributions

AHA contributed to conceptualization, data curation, formal analysis, investigation, methodology, validation, visualization, writing, reviewing and editing; MC contributed to conceptualization, project administration, supervision, review and editing of the manuscript; KP contributed to conceptualization, review and editing of the manuscript; SMR contributed to conceptualization, formal analysis, funding acquisition, methodology, project administration, supervision, visualization, writing, review and editing.

