## Supplemental information for "Developing T cells form an immunological synapse for passage through the β−selection checkpoint"

**This PDF file includes:**

Figures S1 to S9
Tables S1 to S2

### Figure S1

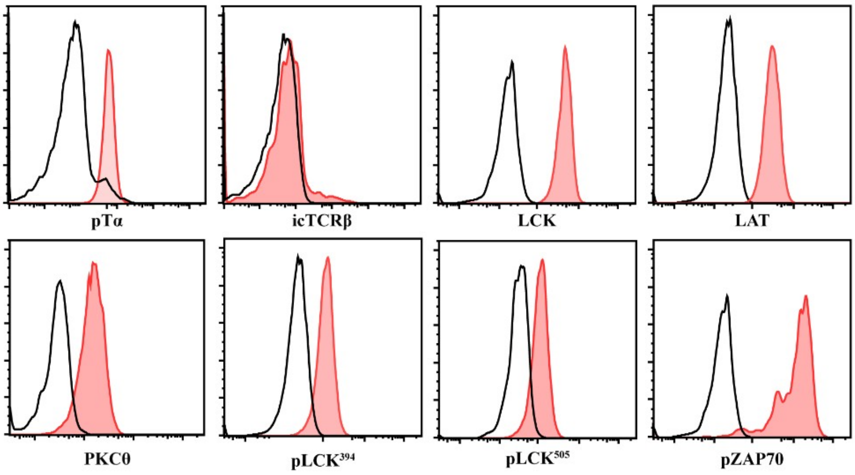

**Fig. S1. Early DN3 cells (DN3a) express pre-TCR structural and signaling molecules.** Primary T cell precursors from mouse fetal liver were co-cultured for 9 days with OP9-DL1 stromal cells. DN3a cells were purified from the co-culture and assessed for the levels of expression and phosphorylation of the pre-TCR structural and signaling molecules as shown by the histograms. The black line indicates Alexa 647 staining without primary antibody, and the red lines are the pre-TCR associated proteins. The results are representative of three independent experiments.

### Figure S2

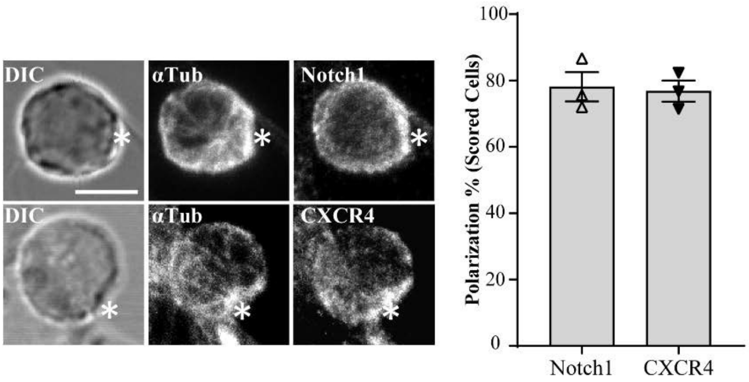

**Fig. S2. Notch1 and CXCR4 polarize at the interface between DN3a and OP9-DL1 stromal cell.** DN3 cells were incubated with OP9-DL1 cells for 14 hours, fixed and stained for α-tubulin to mark the MTOC, and either Notch1 or CXCR4 (as shown). Maximum projection of z-stack images were acquired using confocal microscopy, and representative images shown. After triaging for cells in which MTOC was recruited to the interface (indicated by white asterisk) with an OP9-DL1 cell, the percentage of cells in which Notch1 and CXCR4 were polarized to the interface with the OP9-DL1 cell was determined by blind scoring (as shown by column-bar plot). Total number of scored conjugates per marker is 75, 25 cells per biological replicate. Scale bar, 5µm

### Figure S3

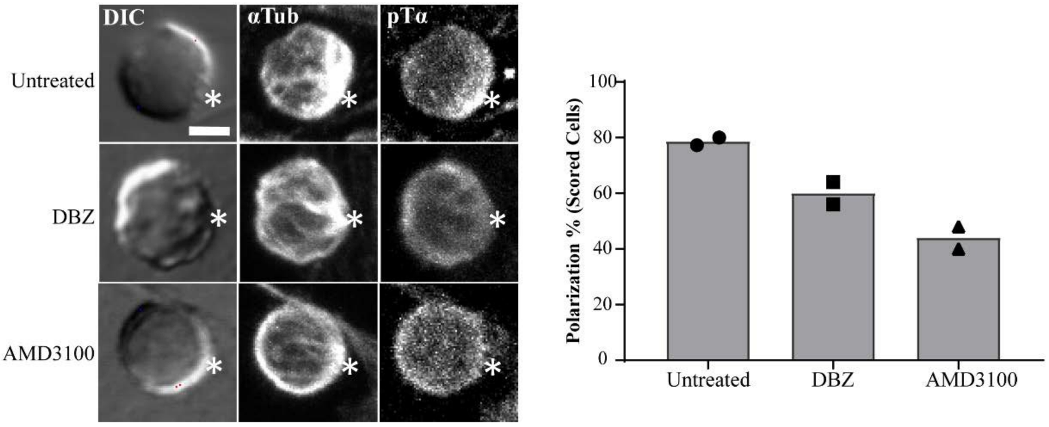

**Fig. S3. Inhibition of Notch and CXCR4 for 3 hours disrupts the immunological synapse formation *in vitro*.** Purified DN3a cells were incubated with OP9-DL1 cells for 3 hours in the presence or absence of either Notch inhibitor (DBZ) or CXCR4 inhibitor (AMD3100), then fixed and stained for α-tubulin to mark the MTOC and pTα chain as a marker of pre-TCR (as shown). After triaging for cells in which MTOC was recruited to the interface with an OP9-DL1 cell, the percentage of cells in which pTα was polarized to the interface with the OP9-DL1 cell was determined by blind scoring (as shown by column-bar plot). Total number of scored conjugates per condition is 50, 25 cells per biological replicate. Scale Bar, 5µm.

Figure S4

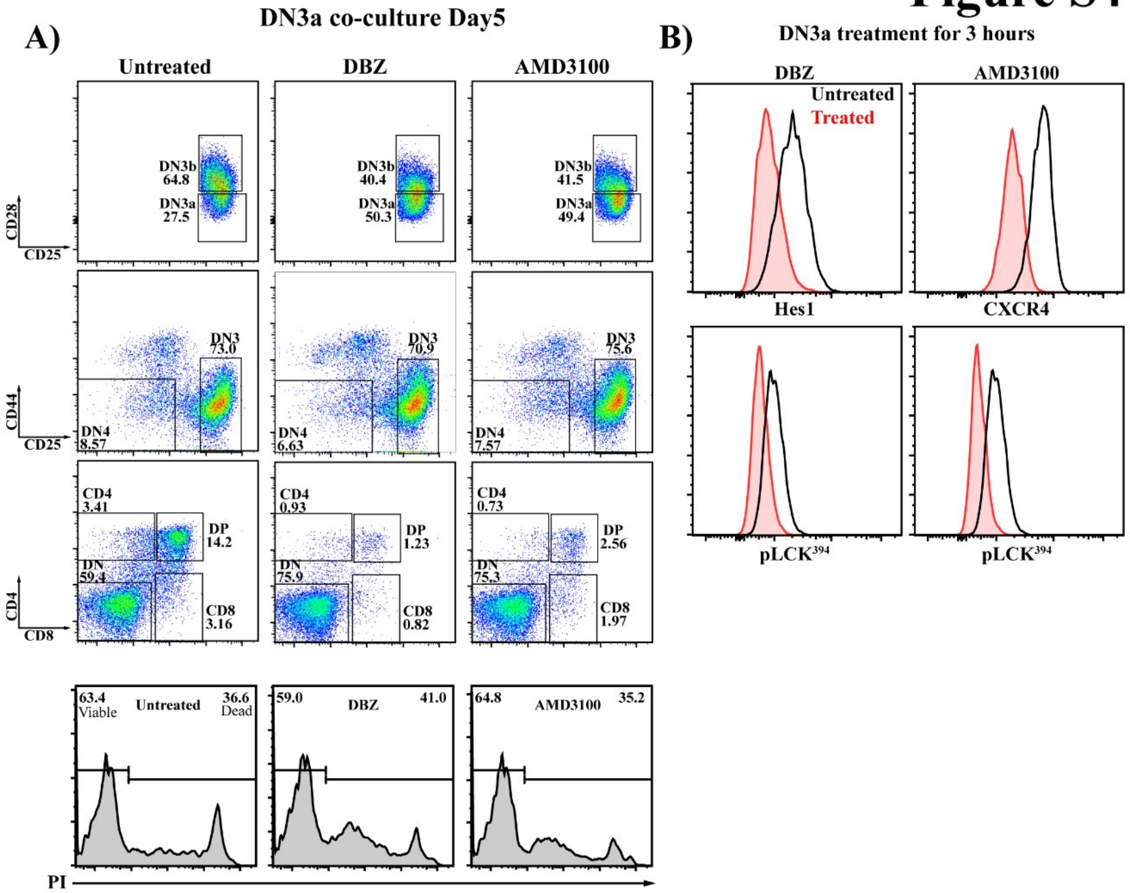

**Fig. S4. Notch and CXCR4 signaling regulate the pre-TCR downstream signaling, and are required for progression beyond β-selection. A)** DN3a cells were co-cultured with OP9-DL1 cells in the presence or absence of either Notch inhibitor (DBZ) or CXCR4 inhibitor (AMD3100). The co-cultures were stopped after 5 days and flow cytometry was used to assess the effect on Notch and CXCR4 inhibition on DN3a cells progression and differentiation (as shown by the flow cytometry plots). The effect of either Notch or CXCR4 inhibition on survival was assessed using PI intensities (as shown by bottom row histograms). **B)** To assess the effect of Notch or CXCR4 inhibition on pre-TCR downstream signaling, purified DN3a co-cultures were treated with either Notch inhibitor (DBZ) or CXCR4 inhibitor for 3 hours, then fixed and levels of $pLCK^{394}$ were assessed as a marker of pre-TCR downstream signaling (as shown by histograms). Hes1 and CXCR4 (top row) were stained to assess the effectiveness of inhibition of Notch and CXCR4, respectively. The results are representative of three independent biological replicates.

### Figure S5

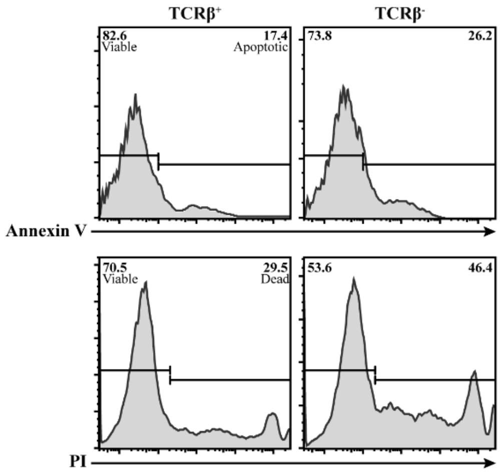

**Fig. S5. DN3a thymocytes expressing surface pre-TCR are more viable that those which does not have pre-TCR expressed.** Purified TCRβ$^+$ DN3a (indicates assembled pre-TCR) and TCRβ$^-$ DN3a (lacks pre-TCR) cells were co-cultured with OP9-DL1 for 5 days. Annexin V dye was used to assess apoptosis, and PI dye was used to assess cell death levels as shown by the histograms. The presented data are representative of three independent biological replicates with similar results.

### Figure S6

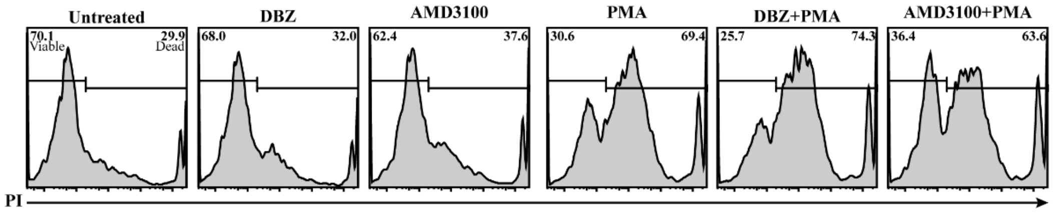

**Fig. S6. DN3a thymocytes differentiating beyond β-selection have poor survival if PMA replaces pre-TCR assembly.** Purified TCRβ$^-$ DN3a cells were co-cultured on OP9-DL1 cells in the presence or absence of either of the following drug combinations; Notch inhibitor (DBZ), CXCR4 inhibitor (AMD3100), a pharmacological mimic of pre-TCR downstream signaling molecule (PMA), PMA with Notch inhibitor and PMA with CXCR4 inhibitor. Co-cultures were stopped after 48 hours and PI was used to assess the effect of each treatment on survival (as shown by histograms). The presented data are representative of three independent biological replicates with similar results.

### Figure S7

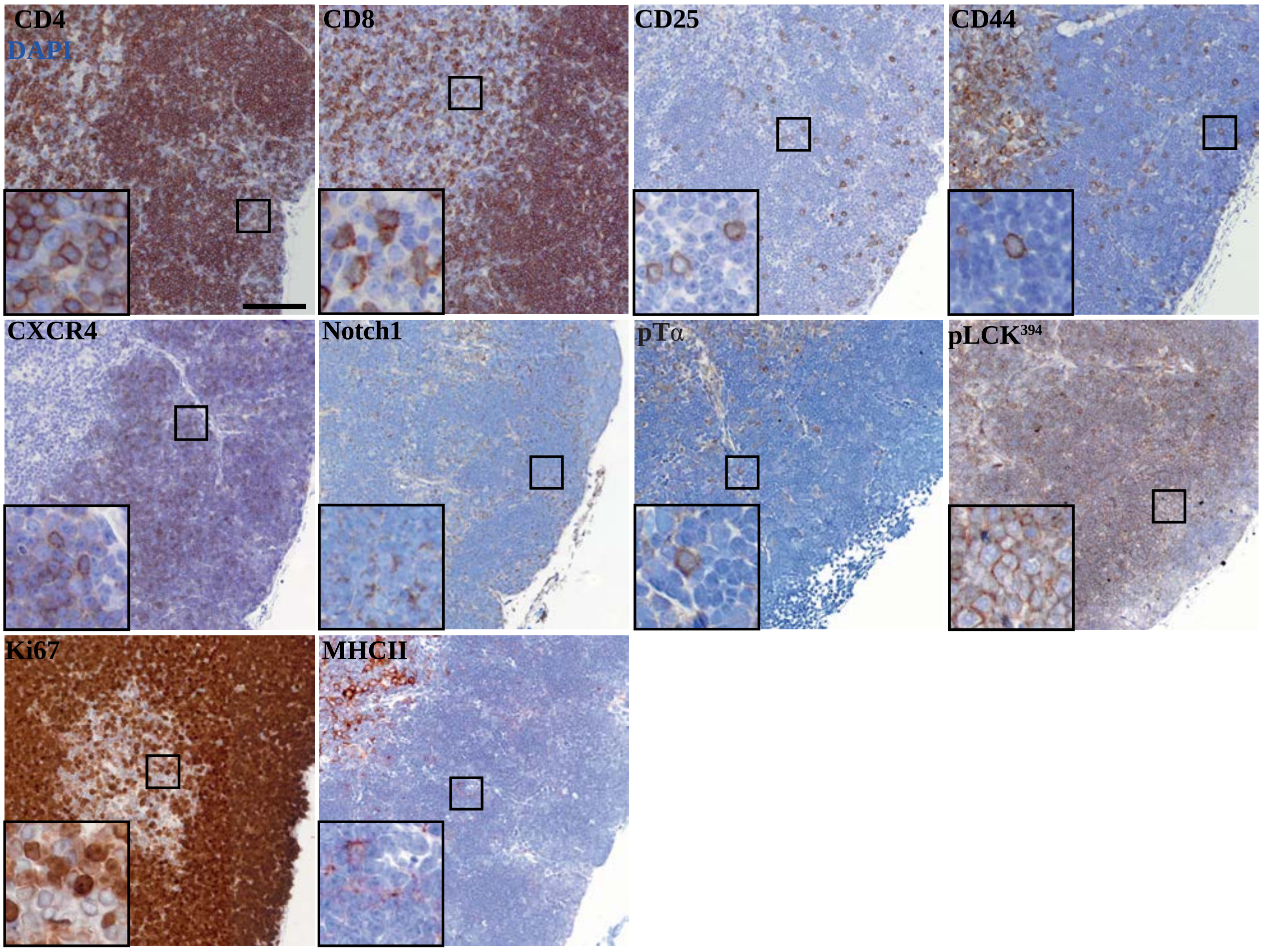

**Fig. S7. Chromogenic detection of markers used in OPAL multiplex.** Thymus lobe slices of 4µm thickness were mounted on glass slide and chromogenic detection (brown) of the multiplex thymus panel markers was carried out, followed counter-stained with hematoxylin to visualize the nucleus (blue) as shown. Images were acquired using Olympus V120 slide scanner and representative images are shown. Scale Bar, 100µm.

### Figure S8

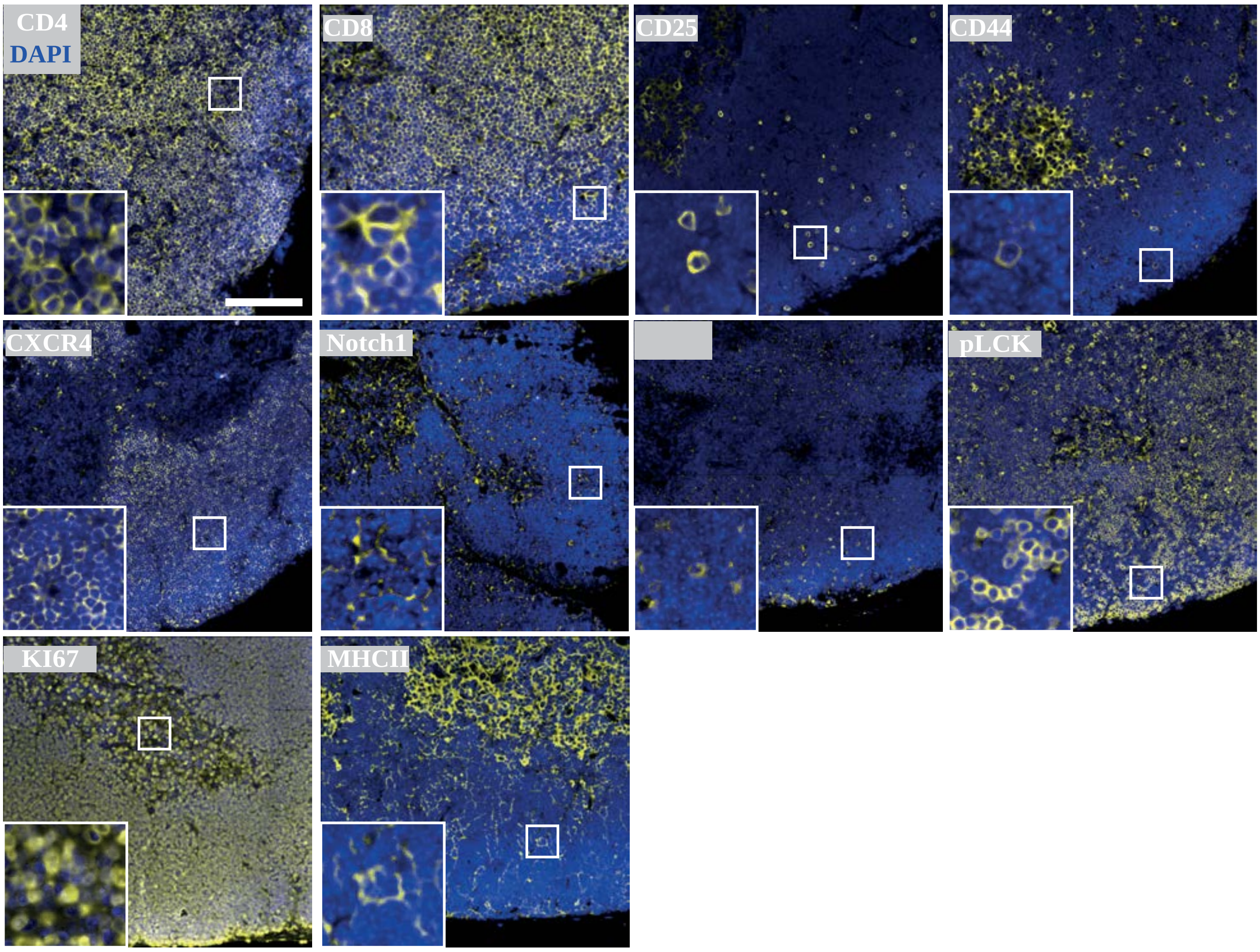

**Fig. S8. Uniplex detection of markers used in OPAL multiplex.** Thymus lobe slices of 4µm thickness were mounted on glass slide and each marker of the multiplex thymus panel was stained with TSA dye (yellow), followed by DAPI staining to mark the nucleus (blue) as shown. Images were acquired using widefield fluorescent microscopy (Vectra® 3 automated quantitative pathology imaging system) and representative images are shown. Scale Bar, 50µm.

### Figure S9

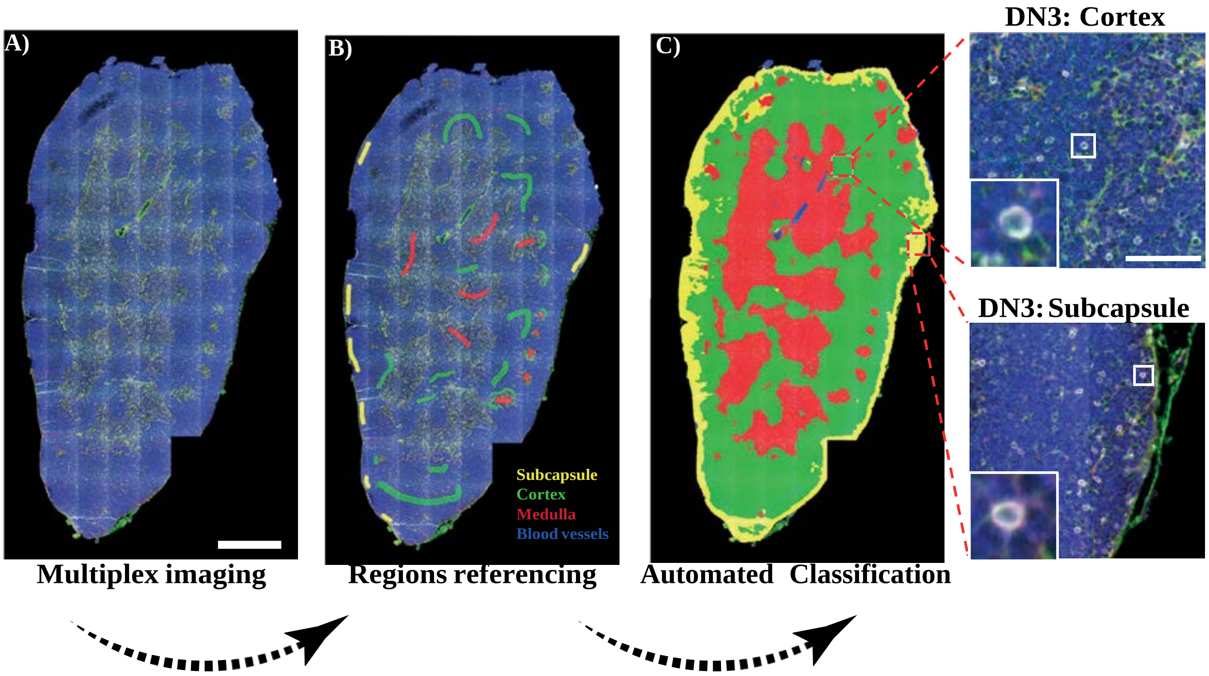

**Fig. S9. Automated region classification of thymic lobes. A)** Thymus lobe slice of 4µm thickness was stained and multiplex imaged using 20X air objective. Images were spectrally dissected then stitched together using HALO indica labs software. **B)** Random regions of the subcapsular (yellow), cortical (green), medullary (red) regions and blood vessels (blue) were marked as reference regions and stored in software library. **C)** An automated classification of entire tissue using reference regions library was carried out. Using CD4, CD8, CD44 and CD25 surface expressions with regions classification we could identify DN3 cells in the cortex (Top zoomed image) or subcapsule (Bottom zoomed image). Scale bar, 1mm (A); 100µm (C, zoomed image).

**Table S1. List of antibodies used in Flow cytometry and immunofluorescence (IF)**

| Antigen | Conjugate | Cat. No | Clone | Reactivity | Host | Supplier | Flow Cytometry (µL) | IF (µL) |
|---|---|---|---|---|---|---|---|---|
| ***CD4*** | PE | 12-0041-81 | RM4-5 | Mouse | Rat | eBioscience | 1:600' | |
| ***CD4*** | Pacific Blue | 48-0042-82 | RM4-6 | Mouse | Rat | eBioscience | 1:1000' | |
| ***CD8α*** | Pacific Blue | 100737 | 53-6.7 | Mouse | Rat | Biolegend | 1:1000' | |
| ***CD44*** | Percp5.5 | 560570 | IM7 | Mouse | Rat | BD Pharmingen | 1:300' | |
| ***CD25*** | BV510 | 102042 | PC61.5 | Mouse | Rat | Biolegend | 1:500' | |
| ***CD28*** | PEcy7 | 122014 | E18 | Mouse | Rat | Biolegend | 1:200' | |
| ***TCRβ*** | APC | 17-5961-82 | H57-597 | Mouse | Armenian Hamster | eBioscience | 1:300' | 1:100' |
| ***pTα*** | | 552407 | 2F5 | Mouse | Mouse | eBioscience | 1:200' | 1:100' |
| ***LCK*** | | MA5-12303 | 3A5 | Mouse | Rabbit | Thermofisher | 1:400' | 1:300' |
| ***pLCK$^{505}$*** | | 2751S | Tyr-505 | Mouse | Rabbit | Cell Signaling | 1:200' | 1:150' |
| ***pLCK$^{394}$*** | | PA537628 | Tyr-394 | Mouse | Rabbit | Thermofisher | 1:250' | 1:150' |
| ***LAT*** | | 22622 | | Mouse | Rabbit | Upstate/Cell signalling | 1:400' | 1:300' |
| ***pZAP70*** | | sc-136248 | pY319.17A | Mouse | Rabbit | Santa Cruz | 1:200' | 1:150' |
| ***PKCϑ*** | | Sc-212 | Poly | Mouse | Rabbit | Santa Cruz | 1:200' | 1:150' |
| ***Notch1*** | | ab27526 | Poly | Mouse | Rabbit | Abcam | 1:400' | 1:200' |
| ***Notch1*** | | AF1057 | Poly | Mouse/Rat | Goat | R&D systems | | 1:200' |
| ***CXCR4*** | | 551968 | 2B11 | Mouse | Rat | BD Pharmingen | 1:300' | 1:300' |

| | | | | | | | |
|---|---|---|---|---|---|---|---|
| ***Alpha-tubulin*** | ab2730 | AP6 | Mouse | Mouse | Abcam | | 1:1000' |
| ***Alpha-tubulin*** | 600-401-880 | | Mouse | Rabbit | Rockland | | 1:500' |
| **Secondary Antibodies** | **Cat. No** | **Conjugate** | **Reactivity** | | **Supplier** | **Flow Cytomtery (µL)** | **IF (µL)** |
| ***Goat α-mouse*** | A-11001 | Alexa 488 | Mouse Primary | | Molecular Probes | 1:1500' | 1:1000' |
| ***Goat α-mouse*** | A-11003 | Alexa 546 | Mouse Primary | | Molecular Probes | 1:1500' | 1:1000' |
| ***Goat α-mouse*** | A-21235 | Alexa 647 | Mouse Primary | | Molecular Probes | 1:1500' | 1:1000' |
| ***Donkey α-goat*** | A-11055 | Alexa 488 | Goat Primary | | Molecular Probes | 1:1500' | 1:1000' |
| ***Donkey α-goat*** | A-21447 | Alexa 647 | Goat Primary | | Molecular Probes | 1:1500' | 1:1000' |
| ***Donkey α-rabbit*** | A-212-6 | Alexa 488 | Rabbit Primary | | Molecular Probes | 1:1500' | 1:1000' |
| ***Donkey α-rabbit*** | A-31572 | Alexa 555 | Rabbit Primary | | Molecular Probes | 1:1500' | 1:1000' |
| ***Donkey α-rabbit*** | A-31573 | Alexa 647 | Rabbit Primary | | Molecular Probes | 1:1500' | 1:1000' |

**Table S2. Thymus Multiplex Antibodies Panel**

| Primary Antigen | Cat. No | Clone | Reactivity | Host | Supplier | OPAL |
|---|---|---|---|---|---|---|
| ***CD4*** | 14-9766-82 | 4SM95 | Mouse | Rat | Thermofisher | 1:1000' |
| ***CD8α*** | 14-0808-80 | 4SM15 | Mouse | Rat | Thermofisher | 1:1000' |
| ***CD44*** | 553132 | IM7 | Mouse | Rat | BD Pharmingen | 1:800' |
| ***CD25*** | PA546922 | Poly | Mouse | Goat | Thermofisher | 1:600' |
| ***pTα*** | 552407 | 2F5 | Mouse | Mouse | eBioscience | 1:200' |
| ***Anti-Mouse*** | PA537628 | Tyr-394 | Mouse | Rabbit | Thermofisher | 1:400' |
| ***Notch1*** | ab27526 | Poly | Mouse | Rabbit | Abcam | 1:400' |
| ***CXCR4*** | 551968 | 2B11 | Mouse | Rat | BD Pharmingen | 1:350' |
| ***Ki67*** | 14-5698-80 | SolA15 | Mouse | Rat | eBioscience | 1:800' |
| ***MHCII*** | 107602 | M5/114.15.2 | Mouse | Rat | Biolegend | 1:500' |

| Secondary Antibodies | Conjugate | Cat. No | Reactivity | Host | Supplier | OPAL |
|---|---|---|---|---|---|---|
| ***Anti-Rabbit*** | HRP | MP-7451 | Rabbit Primary | Horse | Vector Lab. | Ready to use |
| ***Anti-Goat*** | HRP | MP-7405 | Goat Primary | Horse | Vector Lab. | Ready to use |
| | HRP | MP-7452 | Mouse Primary | Goat | Vector Lab. | Ready to use |
| ***Anti-Rat*** | HRP | MP-7404 | Rat Primary | Goat | Vector Lab. | Ready to use |
